# Dopamine D1Aa and D2a Receptor Expression in the Auditory System of a Vocal Fish

**DOI:** 10.1101/2025.11.03.686280

**Authors:** Kobi Kobi, Paul M. Forlano

## Abstract

Although dopamine receptors (DAR’s) have been identified in various vertebrate neural circuits, their expression in the central auditory system remains poorly characterized. Reproductive-state changes in catecholamine innervation of central auditory nuclei in the plainfin midshipman fish (*Porichthys notatus*) highlight a unique, ethologically relevant role for catecholamines, including dopamine, in modulating auditory function to enhance reproductive success. Dopamine’s effects on these systems are mediated via its receptors; thus, the goal of the present study is to characterize excitatory dopamine D1Aa and inhibitory dopamine D2a receptor transcript expression throughout the central auditory system of the plainfin midshipman. Fluorescence in situ hybridization-immunohistochemistry (FISH-IHC) revealed robust D1Aa and D2a expression in forebrain auditory-recipient centers that receive catecholaminergic input: postcommissural and ventral nuclei of the ventral telencephalon, anterior tuberal nucleus, central posterior nucleus of the thalamus, and parvocellular preoptic nuclei. Large dopamine neurons in the periventricular posterior tuberculum, which are, in part, responsible for the reproductive-state changes in central and peripheral catecholamine innervation only express D2a, whereas large noradrenergic neurons in the locus coeruleus express both D1Aa and D2a. The midbrain torus semicircularis, periaqueductal gray, hindbrain octavolateralis efferent nucleus, and descending/secondary octaval nuclei also express both receptor types. D2a expression predominates over D1Aa, and we identify a subpopulation of cells throughout the auditory system that co-express both receptors. The robust distribution of inhibitory and excitatory dopamine receptor expression in the central auditory system, coupled with co-expression in a subset of cells, provides strong neuroanatomical evidence of dopamine’s complex role in modulating auditory sensitivity and processing.

## 1 Introduction

The neurotransmitter dopamine (DA) acts in the central nervous system of vertebrates to modulate a variety of brain functions including motivation, reward, reproduction, motor control, learning, and sensory processing (Forlano, Ghahramani, et al., 2015; Hurley et al., 2004; Kehagia et al., 2010; Matsumoto et al., 1999; Riters, 2012). Among these roles, a growing body of evidence supports a modulatory function for dopamine in auditory sensitivity and processing, with its effects mediated via the activation of specific dopamine receptors (DARs) on target neurons (Gittelman et al., 2013; Missale et al., 1998; Toro et al., 2015). DARs are broadly classified into D1-like receptors, which are generally excitatory and stimulate cAMP production, and D2-like receptors, which are inhibitory and suppress cAMP signaling (Kebabian & Calne, 1979; Missale et al., 1998). In addition to their impact on cAMP production, DARs have also been shown to modulate potassium channels and dopamine transporters, increasing their functional versatility (Liu et al., 1996; Zapata & Shippenberg, 2002). Recent studies have implicated a role for central dopamine receptors modulating auditory processing, including effects on auditory cortex activity and sensory gain control (Happel et al., 2014; Hoyt et al., 2019), and plasticity (Barr et al., 2021; Macedo-Lima et al., 2021). However, these studies have largely focused on candidate brain regions in mammalian and avian systems, and the broader anatomical distribution of dopamine receptors within central auditory circuits, particularly in anamniote vertebrates, remains unclear.

The plainfin midshipman fish (*Porichthys notatus*) is an ideal model for investigating the neural mechanisms underlying auditory function and seasonal plasticity in the auditory system (Bass & McKibben, 2003). During the summer months, male midshipman in the intertidal zone of northern CA and the Pacific Northwest excavate nests under rocks and produce an advertisement call to attract females for mating (Brantley & Bass, 1994). Females must be able to localize these males by sound to lay their eggs and reproduce (Forlano, Sisneros, et al., 2015). Previous work has shown that females undergo a robust enhancement of hearing sensitivity at the level of the inner ear during the summer months to better encode the higher harmonic frequencies of the hum, and thus, better detect the mating call (Sisneros, 2009; Sisneros & Bass, 2003). This seasonal enhancement of auditory sensitivity is, in part, caused by both a summer decrease in dopamine innervation and a decrease in dopamine D2a receptor expression at the level of the inner ear (Forlano, Ghahramani, et al., 2015; Perelmuter et al., 2019). While these studies highlight dopamine’s modulation of hearing sensitivity at the level of the inner ear, robust seasonal differences in catecholamine (CA) innervation, as labeled by fibers immunoreactive for tyrosine hydroxylase (TH; the rate-limiting enzyme in CA synthesis), have also been observed in central auditory nuclei (Forlano, Ghahramani, et al., 2015), presenting an important question of dopamine’s (as well as noradrenaline’s) function in the central auditory system.

Anatomically, the central auditory system in midshipman is very well characterized (see Bass et al., 2000) (Figure 1) with robust CA innervation throughout its extent (Forlano et al., 2014). Dopaminergic neurons in the periventricular posterior tuberculum (TPp) are responsible for the seasonal change in dopamine innervation of the inner ear, send descending projections to the octavolateralis efferent nucleus (OE), descending and secondary octaval nuclei (DO/SO), and ascending projections to the central posterior nucleus of the thalamus (CP), lateral nucleus preglomerulosus (PGl), and some projections to the ventral telencephalon (Vv/Vp/Vs) and parvocellular preoptic area (PPa/PPp) (Forlano et al., 2014; Forlano et al., 2015; Perelmuter & Forlano, 2017). Furthermore, these TPp neurons indirectly receive auditory input from the auditory midbrain (torus semicircularis) via the periaqueductal gray (PAG) (Kittelberger & Bass, 2013), supporting the hypothesis that these neurons and their projections are critically positioned for modulating auditory sensitivity and processing. While the anatomical distribution of CA innervation is well defined, the localization of dopamine receptors within these central auditory regions has not been previously described in midshipman.

**Figure 1:**
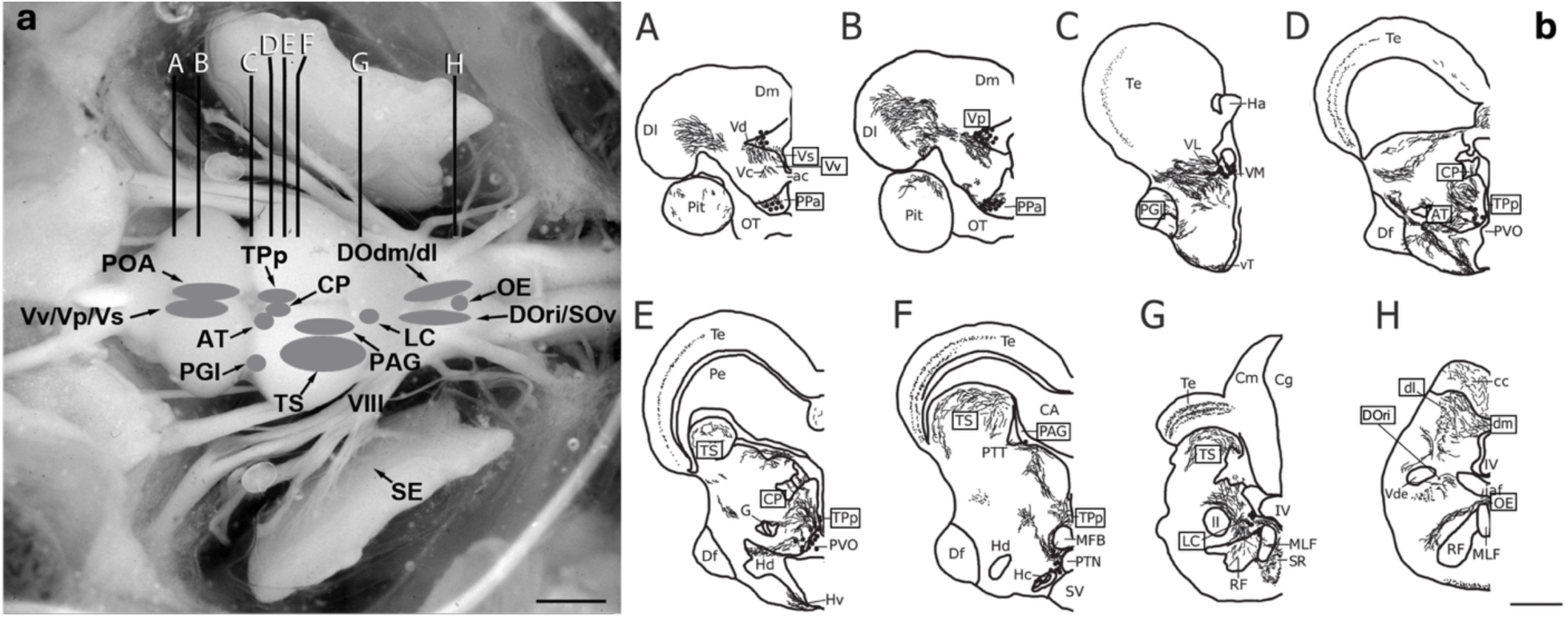
Dorsal view of a midshipman brain (a) with major auditory regions shaded and labeled. Representative series of transverse cross section line drawings (b) illustrating major tyrosine hydroxylase immunoreactive (TH-ir) cell populations, fibers, and terminals. Boxed are auditory and major catecholaminergic regions of interest investigated in the present study. Scale bars = 1.5 mm (a), 500 μm (b). Figures adapted from (Forlano, Ghahramani, et al., 2015). Abbreviations: ac, anterior commissure; AT, anterior tuberal nucleus; CA, cerebral aqueduct; cc, cerebellar crest; Cm, molecular layer of the corpus of the cerebellum; Cg, granular layer of the corpus of the cerebellum; CP, central posterior nucleus of the thalamus; Df, diffuse nucleus of the hypothalamus; dl, dorsolateral division of the descending octaval nucleus; Dl, lateral zone of area dorsalis of the telencephalon; dm, dorsomedial division of the descending octaval nucleus; Dm, medial zone of area dorsalis of the telencephalon; G, nucleus glomerulosus; Ha, habenula; Hc, central periventricular hypothalamus; Hd, dorsal periventricular hypothalamus; Hv, ventral periventricular hypothalamus; iaf, internal arcuate fiber tract; IV, fourth ventricle; LC, locus coeruleus; ll, lateral lemniscus; MFB, medial forebrain bundle; MLF, medial longitudinal fasciculus; OB, olfactory bulb; OE, octavolateralis efferent nucleus; OT, optic tract; PAG, periaqueductal gray; Pe, periventricular cell layer of the torus semicircularis; PPa, anterior parvocellular preoptic nucleus; PGl, lateral division of nucleus preglomerulosus; Pit, pituitary; POA, preoptic area; PTN, posterior tuberal nucleus; PTT, paratoral tegmentum; PVO, paraventricular organ; RF, reticular formation; SE, saccular epithelium; SR, superior raphe; SV, saccus vasculosus; Te, midbrain tectum; TPp, periventricular posterior tuberculum; TS, torus semicircularis; Vc, central nucleus of area ventralis of the telencephalon; Vd, dorsal nucleus of area ventralis of the telencephalon; Vde, descending tract of the trigeminal nerve; VIII, auditory eighth nerve; VL, ventrolateral nucleus of the thalamus; VM, ventromedial nucleus of the thalamus; Vp, postcommissural nucleus of area ventralis of the telencephalon; Vs, supracommissural nucleus of area ventralis of the telencephalon; vT, ventral tuberal hypothalamus.

To begin to address dopamine’s functional impact on the central auditory system, we compared the expression of an excitatory dopamine receptor, D1Aa, and an inhibitory dopamine receptor, D2a, in central auditory nuclei in the brain of adult midshipman fish. These dopamine receptors were two of seven that were previously identified in the midshipman inner ear (Perelmuter et al., 2019). The sequence of the D1A receptor gene is the most conserved in vertebrates, and the D1Aa subtype in another teleost fish, the zebrafish, is the only D1A subtype that shows synteny shared with other species, therefore, showing greater conservation of this gene (Yamamoto et al., 2015). Thus, of the excitatory D1-like class of receptors, the D1Aa receptor gene is an ideal candidate for characterization in the central auditory system. D2 receptors are also highly conserved compared to other D2-like receptors (D3, D4) and exhibit a close phylogenetic relationship with other vertebrate D2 receptors in another teleost fish, the European eel (Pasqualini et al., 2009). Furthermore, because expression of the inhibitory D2a receptor is seasonally regulated in the inner ear (Perelmuter et al., 2019), the D2a receptor gene is an important candidate for characterization in the central auditory system. Here, we show cells in various central auditory centers that are both innervated by catecholaminergic fibers and express D1Aa and/or D2a receptor transcripts.

## 2 Materials and Methods

### 2.1 Animals

Female plainfin midshipman fish in reproductive condition (n=12) were collected at low tide by hand in the rocky intertidal zones of Hood Canal, Washington and shipped to the Aquatic Research and Environmental Assessment Center (AREAC) at City University of New York (CUNY) Brooklyn College. Animals were briefly maintained in recirculating saltwater aquaria between 1-5 days before sacrifice. All acquisition, maintenance, and experimental procedures on animals performed in this study were approved by the Institutional Animal Care and Use Committee (IACUC) of CUNY Brooklyn College.

### 2.2 Tissue Collection and Preparation

Animals were anesthetized in 0.025% benzocaine solution in seawater and sacrificed via a transcardial perfusion of teleost ringers (4°C) followed by 4% paraformaldehyde (PFA) in phosphate buffer (0.1M PB, pH = 7.2, 4°C). Brains were dissected, postfixed in PFA for 1 hour, rinsed with PB, and cryoprotected in 30% sucrose-PB solution for 48 hours at 4°C. Brains were then embedded in Cryo-Gel (Leica Surgipath), sectioned at 20 µm in the transverse plane on a Leica CM1850 cryostat, collected on positively charged microscope slides (Globe Scientific), and stored at -80°C until fluorescence in situ hybridization.

### 2.3 Hybridization Probes and Amplification Hairpins

Sequences for the plainfin midshipman dopamine D2a and dopamine D1Aa receptor that were previously identified (Perelmuter et al., 2019) were sent to Molecular Instruments (Los Angeles, California). Hybridization probes with split-initiators (B1 and B2) targeting the specific transcripts were designed and manufactured, and B1 and B2 amplification hairpins were also purchased from Molecular Instruments. Hybridization probes and amplification hairpins used in this study are listed in Table 1.

**Table 1:** Hybridization Probes and Amplification Hairpins.

| Receptor Gene | GenBank Accession # | Initiator/Amplifier-Fluorescent Dye | Lot Number |
| --- | --- | --- | --- |
| Dopamine D1Aa | GBYQ01030481 | B1 Alexa Fluor 488 | S020925 |
| Dopamine D2a | GBYQ01052780 | B2 Alexa Fluor 594 | S017125 |

### 2.4 Fluorescence in Situ Hybridization

Slides were removed from -80°C and immediately fixed in 4% PFA (in 0.1M PB, 4°C) for 15 minutes followed by a dehydration series of ethanol (50%/70%/100%/100%) for 5 minutes each. Slides were briefly soaked in phosphate buffered saline (0.1M PBS, pH = 7.2) before a hydrophobic border was applied (Vector Laboratories ImmEdge Pen). Sections were soaked 2 x 15 minutes with a solution of 3% hydrogen peroxide (3% H2O2, 20 mM NaOH, 0.1M PBS), washed with PBS 3 x 5 minutes, and incubated in probe hybridization buffer (Molecular Instruments) for 10 minutes at 37°C. Probe hybridization solution (16 nM D1Aa and 8 nM D2a in pre-heated (37°C) probe hybridization buffer) was prepared and applied to slides before an overnight incubation (∼20 hours) at 37°C. Specific probe concentrations were optimized to achieve the highest signal to noise. After the HCR RNA-FISH hybridization, slides were washed with a series of probe wash buffer (PWB, Molecular Instruments)/saline-sodium citrate tween buffer (SSCT) at 37°C for 15 minutes each: 100% PWB/75% PWB/50% PWB/25% PWB. Slides were then washed in 100% SSCT for 15 minutes at room temperature and treated with amplification buffer (Molecular Instruments) for 1 hour. Amplification solution was prepared by heating 6 pmol of hairpin h1 and h2 for both B1 and B2 at 95°C for 90 seconds, cooling in a dark drawer for 30 minutes, and then combining all hairpins with amplification buffer. Amplification solution was applied to slides before an overnight incubation (∼20 hours) at room temperature. Slides were washed 3 x 30 minutes with SSCT before proceeding to fluorescence immunohistochemistry.

### 2.5 Fluorescence Immunohistochemistry

Slides were washed 3 x 10 minutes in PBS and then blocked for 1 hour in PBST (0.1 M PBS, 5% normal donkey serum (NDS, Jackson ImmunoResearch), 0.3% Triton X-100). Primary antibody solution was prepared as in previous studies (Forlano et al., 2014) with mouse anti-tyrosine hydroxylase (TH, 1:1000 dilution in PBST, EMD Millipore/Chemicon MAB318, RRID: AB_2201528, Immunogen: Tyrosine Hydroxylase purified from PC12 cells, clone LNC1). Primary antibody solution was applied to tissue sections and incubated overnight (∼20 hours) at room temperature in a humidified chamber. Following, slides were washed 5 x 10 minutes in 0.1M PBS + 0.5% NDS and incubated with Alexa Fluor 680 diluted in PBST (1:200, anti-mouse Alexa Fluor 680, Invitrogen) for 2 hours. Finally, slides were washed 4 x 10 minutes in PBS and cover slipped with Prolong Gold Antifade Reagent with 4,6-diamidino-2-phenylindole (DAPI) nuclear counterstain (Invitrogen).

### 2.6 Fluorescent Image Acquisition

All images were taken on an Olympus BX61 epifluorescence compound microscope using various objectives (20x, 40x, 60x) with DAPI, GFP, Texas Red, and Cy5.5 (ultra far red) filters (Chroma). Images were acquired with Metamorph imaging software (Molecular Devices) as 5-11 µm z-stacks with 0.2-1 µm steps depending on the objective. Exposure time, number of z-stacks, and step size were all optimized for each image and region of interest. Photomicrographs were compiled via a maximum intensity projection of each channel, contrast-enhanced on Metamorph or ImageJ, and overlaid into a single image.

## 3 Results

### 3.1 FISH-IHC Overview

An anatomical overview of all auditory regions sampled is indicated in Fig 1. A summary of D1Aa and D2a expression with relative abundance in central auditory regions is provided in Table 2, which also highlights the expression of androgen and estrogen receptors, as well as aromatase, which converts testosterone to estrogen (Fergus & Bass, 2013; Forlano et al., 2001, 2005, 2010; Forlano & Sisneros, 2016).

**Table 2:** Comparative distribution of dopamine D1Aa, dopamine D2a, androgen receptor (AR), aromatase (ARO), estrogen receptor alpha (ERα), estrogen receptor beta 1 (ERβ1), and estrogen receptor beta 2 (ERβ2) in central auditory centers and other areas of interest.

| Brain Area | D1Aa | D2a | AR | ARO | ERα | ERβ1 | ERβ2 |
| --- | --- | --- | --- | --- | --- | --- | --- |
| **Forebrain** |  |  |  |  |  |  |  |
| Postcommissural Nucleus of the Ventral Telencephalon (Vp) | ++ | ++ | ++ | +++ | + | + | - |
| Ventral Nucleus of the Ventral Telencephalon (Vv) | ++ | ++ | - | +++ | + | - | + |
| Supracommissural Nucleus of the Ventral Telencephalon (Vs) | (-) | +++ | +++ | +++ | + | - | + |
| Anterior Parvocellular Preoptic Nucleus (PPa) | + | ++ | +++ | +++ | +++ | + | + |
| Posterior Parvocellular Preoptic Nucleus (PPp) | ++ | ++ | ++ | +++ | - | - | + |
| **Midbrain/Diencephalon** |  |  |  |  |  |  |  |
| Periventricular Posterior Tuberculum (TPp) | - | +++ | ++ | +++ | + | - | + |
| Torus Semicircularis (TS) | ++ | ++ | ++ | - | - | - | + |
| Periaqueductal Gray (PAG) | ++ | +++ | + | + | - | + | + |
| Anterior Tuberal Nucleus (AT) | + | + | ++ | +++ | +++ | - | + |
| Central Posterior Nucleus of the Thalamus (CP) | + | ++ | + | +++ | + | - | - |
| Lateral Division of the Nucleus Preglomerulosus (PGL) | (-) | ++ | + | + | - | - | - |
| **Hindbrain** |  |  |  |  |  |  |  |
| Octavolateralis Efferent Nucleus (OE) | ++ | ++ | - | - | - | - | - |
| Dorsomedial/Dorsolateral Division of the Descending Octaval Nucleus (DOdm/dl) | + | ++ | + | - | - | - | - |
| Rostral Intermediate Division of the Descending Octaval Nucleus (DOri) | + | ++ | - | - | - | - | - |
| Ventral Division of the Secondary Octaval Nucleus (SOv) | + | +++ | - | - | - | - | - |
| Locus Coeruleus (LC) | ++ | +++ | - | - | - | - | - |

‘+++’, ‘++’, ‘+’ represent relative levels of hybridization signal of D1Aa, D2a, AR, Aro, and ERα . Expression of ERβ1 and ERβ2 is represented by ‘+’ as relative levels of hybridization signal are unknown. ‘-’ represents zero signal whereas ‘(-)’ represents signal that is inconclusively above background. Table modified from Forlano et al., (2010), Fergus and Bass (2013), Forlano et al., (2016).

### 3.2 Telencephalon

Robust labeling of D1Aa and D2a occurs throughout the area ventralis (V) and preoptic area (PPa/PPp). While these regions are not exclusively auditory in function, they receive input projections from auditory nuclei and the TPp (Bass et al., 2000; Forlano et al., 2014; Goodson & Bass, 2002) and therefore included in this study. Within the postcomissural (Vp) and ventral nuclei (Vv), many cells co-express D1Aa and D2a (Figs, 2A-F). DAR expression is largely uniform throughout the entirety of both nuclei. D2a seems to predominate over D1Aa within cells that co-express D1Aa/D2a, and there is a population of cells in both nuclei that express D2a but not D1Aa (Figs 2A-F). Within the supracomissural nucleus (Vs), cells express D2a (Fig 2G); however, D1Aa signal is not above background (not shown). All of these regions are potential targets of the local TH-ir cell population (shown in Fig 2A), which also express D2a, and the TPp (Forlano et al., 2014; Northcutt, 2006; Tay et al., 2011). Furthermore, within the anterior and posterior parvocellular preoptic nuclei (PPa/PPp), many cells co-express D1Aa and D2a (Figs 3A-F) with a uniform robust expression throughout the nuclei.

**Figure 2:**
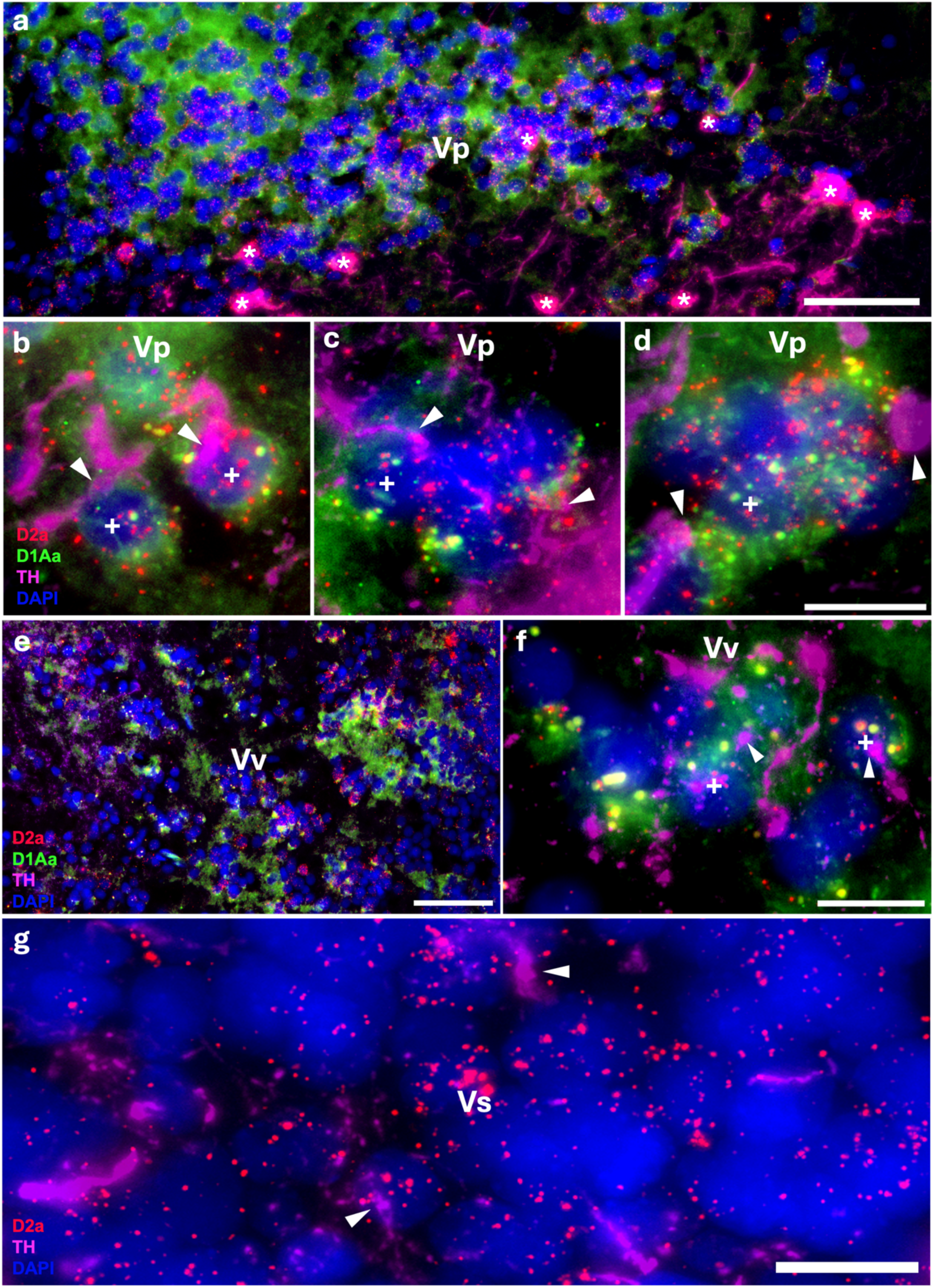
Robust D1Aa (green)/D2a (red) expression and TH-ir (magenta) innervation of forebrain nuclei; nuclei stained with DAPI in blue. **A-F:** TH-ir fibers innervating/terminating at postcomissural nucleus of area ventralis (Vp) cells **(A-D)** and ventral nucleus of area ventralis (Vv) cells **(E, F)** co-expressing D1Aa and D2a receptor transcripts (labeled with +). TH-ir cell population lies just ventral to the postcomissural nucleus of area ventralis (Vp, A, * denotes TH-ir cell). **G:** TH-ir fibers innervating/terminating at supracomissural nucleus of area ventralis (Vs) cells expressing D2a receptor transcripts. All arrowheads point to TH-ir fibers/putative terminals. Scale bars in A, E = 50 μm; B, C, D, F, G = 10 μm.

**Figure 3:**
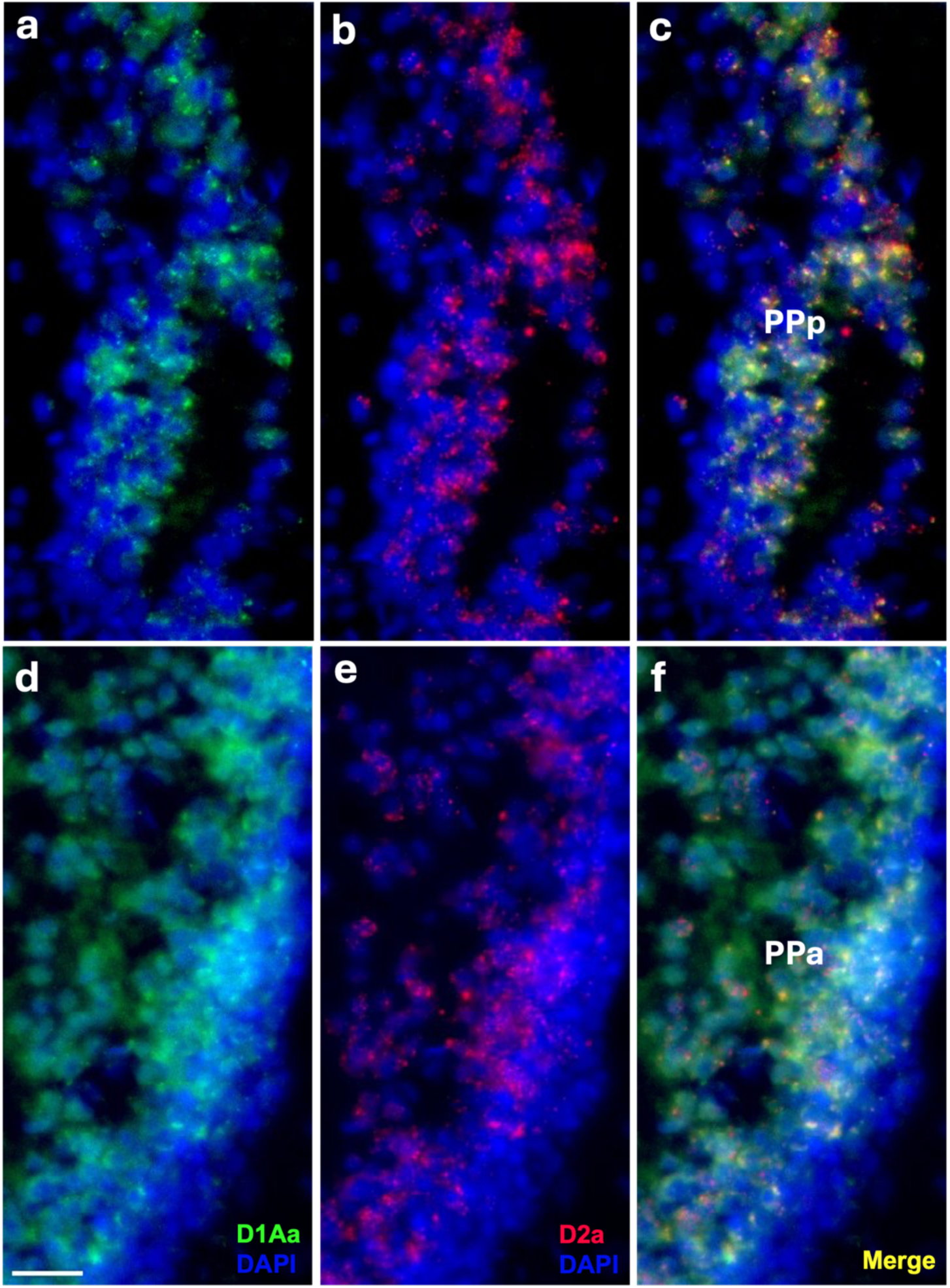
Robust D1Aa (green)/D2a (red) expression in the posterior parvocellular preoptic area (PPp, **A-C)** and the anterior parvocellular preoptic area (PPa, **D-F)**; nuclei stained with DAPI in blue. Scale bar = 20 μm.

### 3.3 Periventricular Posterior Tuberculum

There is robust D2a expression in the large TH-ir cells of the periventricular posterior tuberculum (TPp, Figs 4A-B), which provide dopaminergic efferents to the peripheral auditory system as well as all levels of the central auditory system (Forlano et al., 2014; Perelmuter & Forlano, 2017). Within the entire brain, D2a expression is perhaps the most robust in these dopaminergic TPp cells and is one population that we identified to conclusively not express D1Aa. While the TPp broadly has high D2a expression relative to other regions, there is some variation of expression within each individual TH-ir TPp cell. Despite this, the expression pattern and variation seem uniform throughout the rostral-caudal length of the nucleus.

**Figure 4:**
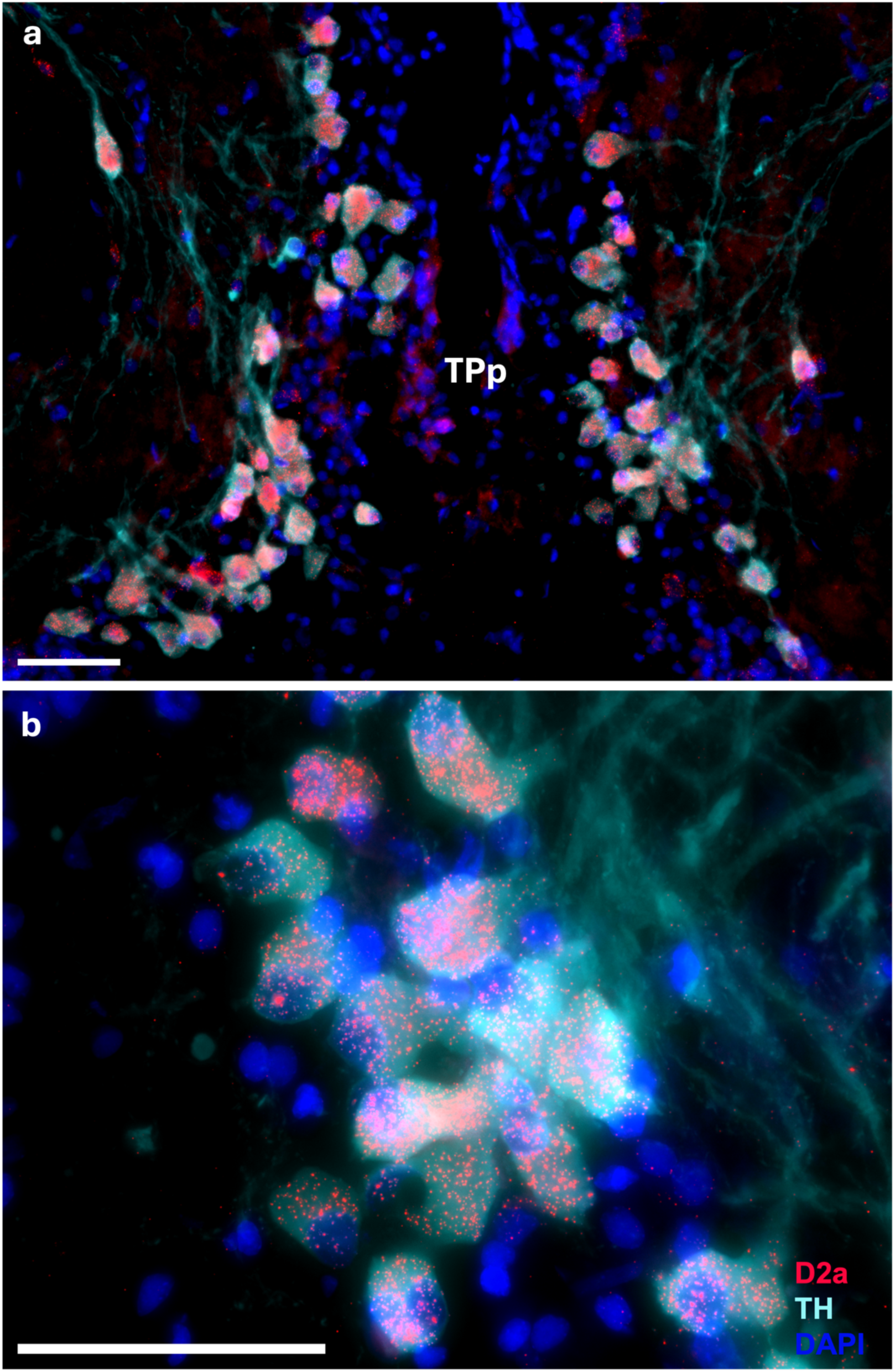
Robust D2a expression (red) on somata of large dopamine neurons labeled by TH-ir (cyan) in the periventricular posterior tuberculum (TPp); nuclei stained with DAPI in blue. Scale bars = 50 μm.

### 3.4 Auditory Diencephalon and Midbrain

Robust labeling of D1Aa and D2a occurs throughout the auditory midbrain/diencephalon. Within the torus semicircularis nucleus centralis (TSnc), the main central auditory relay center to higher order nuclei (Bass et al., 2000), and the periaqueductal gray (PAG), an auditory-vocal integration site with reciprocal connections with the TPp (Kittelberger & Bass, 2013), there are a subset of cells that co-express D1Aa and D2a, and another subset of cells that only express D2a (Figs. 5A-G). Within the anterior tuberal nucleus (AT) and central posterior nucleus of the thalamus (CP), auditory regions that receive input from both the TS and TPp (Bass et al., 2000; Forlano et al., 2014), there also seems to be a subset of cells that co-express D1Aa and D2a, and another subset of cells that only express D2a (Figs. 6A-J). Furthermore, TH-ir cells in the AT seem to show a very robust expression of D2a (Fig 6A). In the nucleus preglomerulosus (PGl), which also receives input from the TS, there is a subset of cells that express D2a (Fig 6K). However, D1Aa signal is not conclusively above background (not shown). Interestingly, in all these regions, there seems to be great variation of receptor expression within cells (see arrowheads in Fig 6K). Despite this, the expression pattern and variation seem uniform throughout the rostral-caudal length of these nuclei.

**Figure 5:**
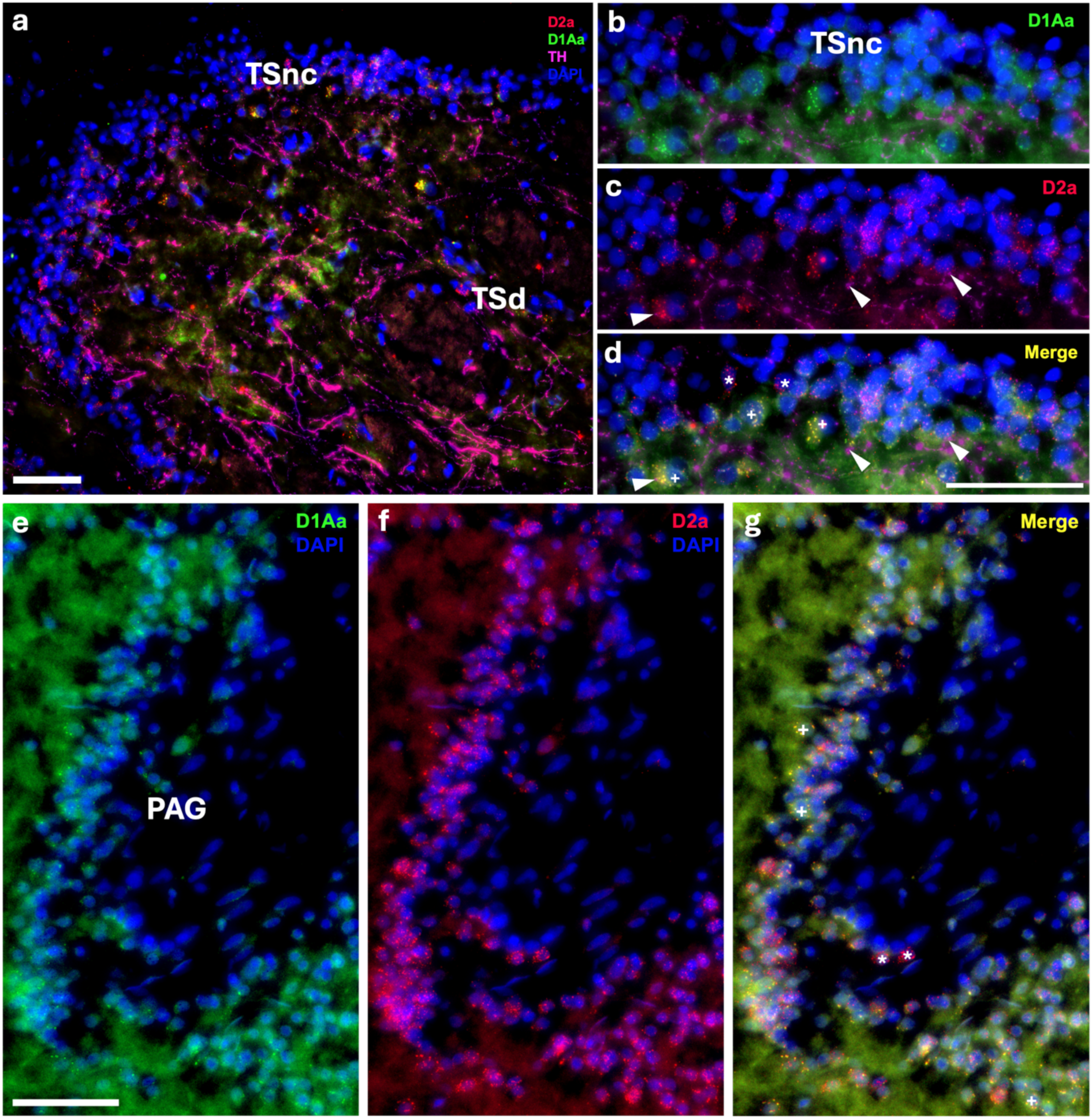
Robust D1Aa (green)/D2a (red) expression and TH-ir (magenta) innervation of midbrain nuclei stained with DAPI in blue. **A-D:** TH-ir fibers innervating/terminating at torus semicircularis nucleus centralis (TSnc) cells co-expressing D1Aa and D2a. **E-G:** Robust D1Aa/D2a expression in periaqueductal gray (PAG) cells. + labeled cells are D1Aa+/D2a+ and * labeled cells are D1Aa-/D2a+. Scale bars in A, E, F, G = 50 μm; B, C, D = 10 μm.

**Figure 6:**
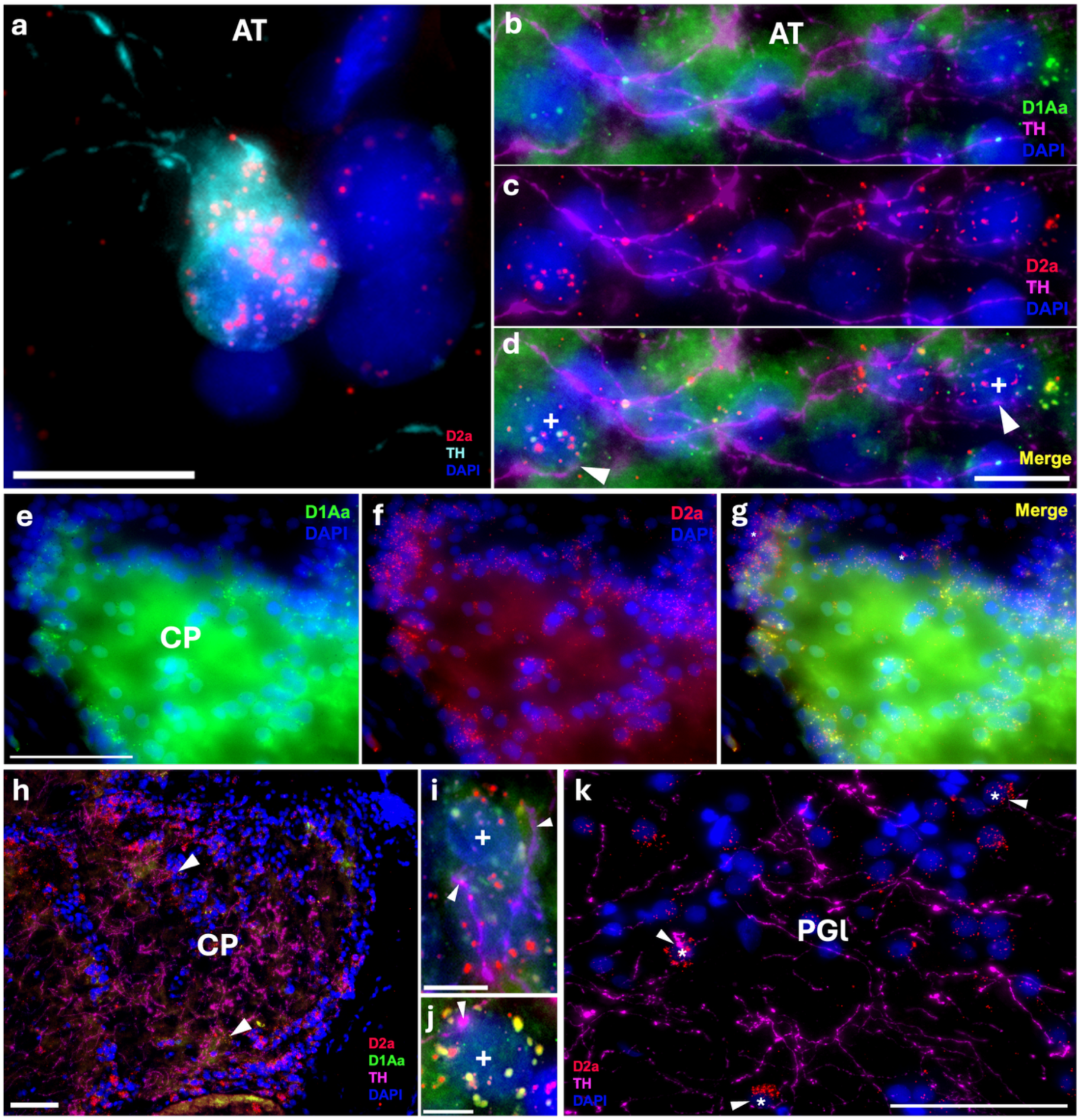
Robust D1Aa (green)/D2a (red) expression and TH-ir (magenta) innervation of diencephalic/midbrain auditory centers; nuclei stained with DAPI in blue. **A:** TH-ir cell (cyan) in the anterior tuberal nucleus (AT) expressing D2a (red). **B-D:** Other cells in the AT surrounded by TH-ir fibers (magenta) and co-expressing D1Aa and D2a. **E-G:** Cells in the central posterior nucleus of the thalamus (CP) co-expressing D1Aa and D2a. **H-J:** TH-ir fibers innervating/terminating at CP cells co-expressing D1Aa and D2a. **K:** TH-ir fibers innervating/terminating at lateral nucleus preglomerulosus (PGl) cells expressing D2a. + labeled cells are D1Aa+/D2a+ and * labeled cells are D1Aa-/D2a+. All arrowheads point to TH-ir fibers/putative terminals; arrowheads in H point to TH-ir fibers/putative terminals on cells that are shown in I and J. Scale bars in A, B, C, D = 10 μm; E, F, G, H, K = 50 μm; I, J = 5 μm.

### 3.5 Hindbrain

In the octavolateralis efferent nucleus (OE), which sends cholinergic projections to the inner ear (Bass et al., 1994; Highstein & Baker, 1986) and receives its sole catecholaminergic input from dopamine originating from the TPp (Perelmuter & Forlano, 2017), a subset of cells appear to co-express D1Aa and D2a (Figs 7A-B). This is consistent in both the caudal (OEc) and rostral (OEr) subdivisions. In the locus coeruleus (LC), which is the primary source of noradrenaline and receives dopaminergic input from the TPp (Kaslin & Panula, 2001) all cells seem to co-express D1Aa and D2a (Figs 7C-E), which is consistent throughout the rostral-caudal length of the LC.

**Figure 7:**
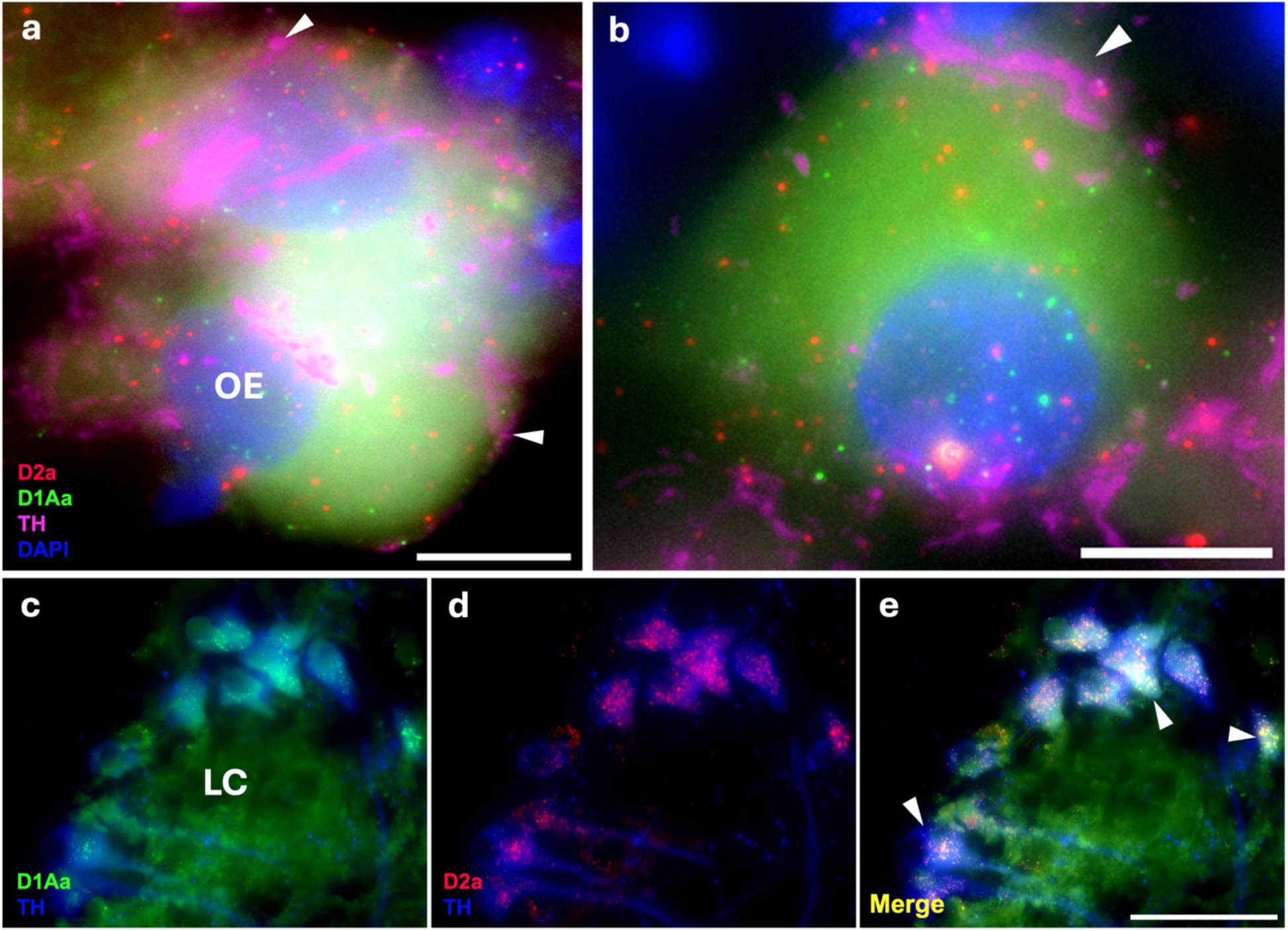
D1Aa (green)/D2a (red) expression and TH-ir (magenta) innervation in hindbrain. **A-B:** TH-ir fibers innervating/terminating at octavolateralis efferent nucelus (OE) cells co-expressing D1Aa and D2a. Arrowheads point to TH-ir fibers/putative terminals **C-E:** Robust D1Aa/D2a expression in the noradrenergic locus coeruleus (LC). Arrowheads point to TH-ir LC cells in blue. Scale bars in A, B = 10 μm; C, D, E = 50 μm.

At all three levels of the descending octaval nucleus (dorsomedial-DOdm, Figs 8A-C; dorsolateral-DOdl, Figs 8D-F; and rostral intermediate-DOri, Figs 8G-I), as well as the secondary octaval nucleus (SOv, Figs 8J-L), there is a subset of cells with robust co-expression of D2a and D1Aa. Both the DO and the SO receive input from primary auditory afferents and are densely innervated by TH-ir fibers, likely originating from the TPp (Bass et al., 2001; Forlano et al., 2014). All these regions also contained a subset of cells that only expressed D2a, and we did not identify any cells that solely expressed D1Aa. Within the three levels of the DO, the expression pattern was consistent throughout the nuclei proper. Within the SOv, expression of both receptors seemed to increase with the higher TH-ir innervation in the rostral part of the nucleus.

**Figure 8:**
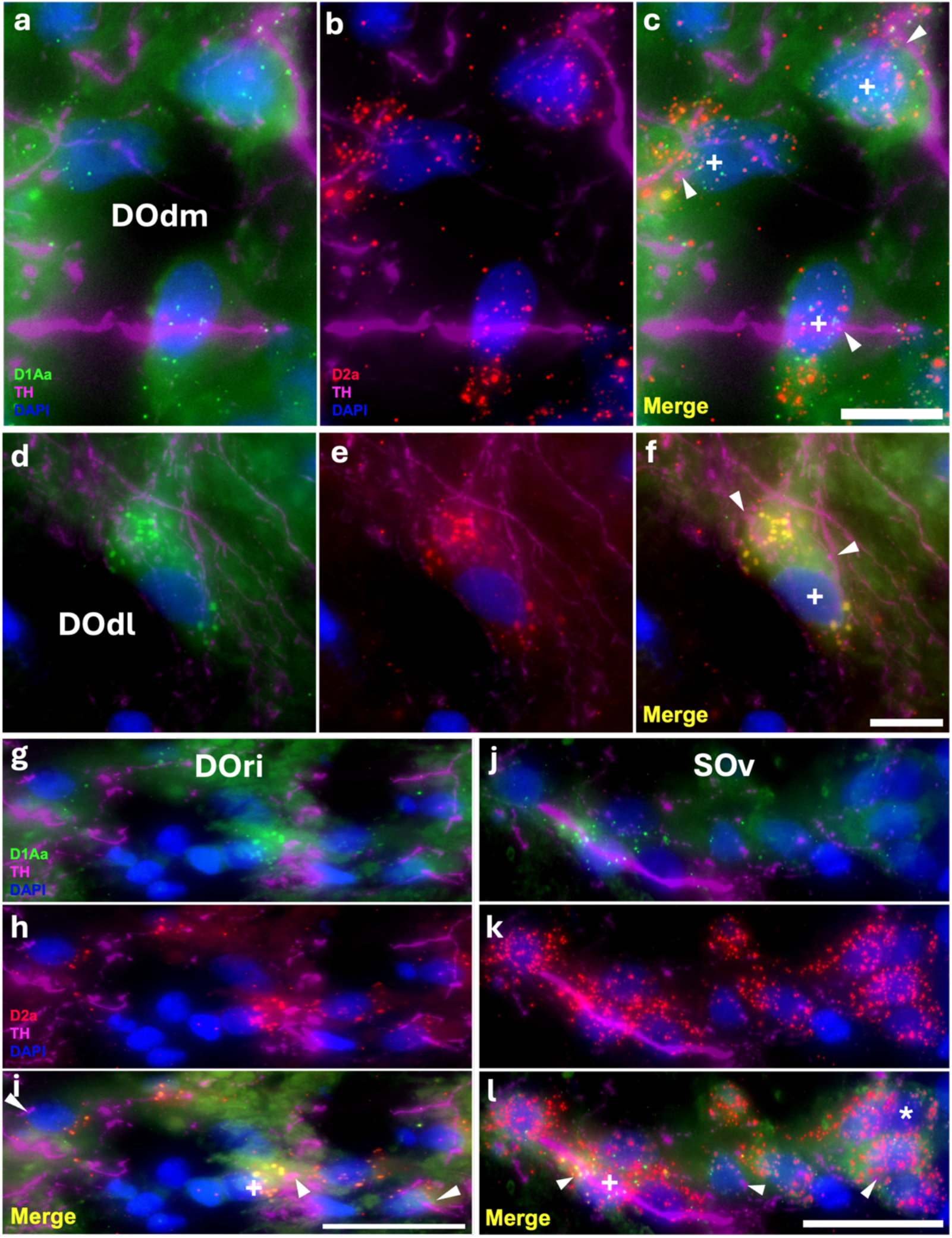
Robust D1Aa (green)/D2a (red) expression and TH-ir (magenta) innervation of hindbrain auditory nuclei; nuclei stained with DAPI in blue. **A-I:** TH-ir fibers innervating/terminating at dorsomedial (DOdm, **A-C**), dorsolateral (DOdl, **D-F**), and rostral intermediate (DOri, **G-I**) cells of the descending octaval nucleus (DO) co-expressing D1Aa and D2a. **J-L:** TH-ir fibers innervating/terminating at ventral secondary octaval nucleus (SOv) cells expressing D1Aa and D2a. + labeled cells are D1Aa+/D2a+ and * labeled cells are D1Aa-/D2a+. All arrowheads point to TH-ir fibers/putative terminals. Scale bars in A-F = 10 μm; G-L = 25 μm.

## 4 Discussion

### 4.1 Overview

In this study, we characterized dopamine D1Aa and D2a receptor transcript mRNA distribution throughout the central auditory system of the plainfin midshipman fish. To our knowledge, this is the first study to broadly characterize both excitatory and inhibitory dopamine receptor expression together in an auditory system of a vertebrate as well as demonstrate robust co-expression of excitatory and inhibitory dopamine receptors within individual neurons spanning hindbrain to forebrain. Furthermore, by combining FISH with traditional IHC to label for tyrosine hydroxylase (TH), we provide strong neuroanatomical support for dopamine innervation and modulation in specific auditory nuclei via its receptor expression. While TH is a marker for both dopaminergic and noradrenergic cells and fibers, our previous studies have demonstrated that TPp neurons, known to be dopaminergic (see below), project specifically to several auditory nuclei, and the auditory hindbrain is largely devoid of noradrenergic fibers (Forlano et al., 2014; Perelmuter & Forlano, 2017). Although we focused primarily on two dopamine receptors in the central auditory system (see section 1), future studies characterizing the central expression of the other five dopamine receptors in midshipman are needed to better understand how dopamine modulates auditory sensitivity and processing, as well as its role in neuroendocrine, motivational, and motor functions. Furthermore, a quantitative/seasonal analysis of dopamine receptor expression was beyond the scope of this initial study and will be reported elsewhere.

### 4.2 D2a Expression in the Periventricular Posterior Tuberculum

The TH-ir neurons in the periventricular posterior tuberculum (TPp) show the most robust expression of D2a throughout the entire brain. Many studies confirm that these cells are indeed DAergic as they are immunoreactive to DA and TH but not dopamine β-hydroxylase and contain genetic markers specific for dopamine neurons in mammals (Batten et al., 1993; Ekström et al., 1986; Filippi et al., 2010; Forlano et al., 2014; Kaslin & Panula, 2001; Perelmuter & Forlano, 2017; Ryu et al., 2007; Yamamoto et al., 2011). Furthermore, previous work has shown that these TPp neurons send strong descending projections to the hindbrain, local projections within the diencephalon, and ascending projections to the ventral telencephalon-all regions that were investigated in this study (Forlano et al., 2014; Ma, 2003; Tay et al., 2011). Seasonal changes in CA innervation of various central auditory centers as well as the inner ear are either entirely or in part caused by the TPp (Forlano, Ghahramani, et al., 2015; Perelmuter et al., 2021), thus strongly implicating an important role for these dopaminergic cells in modulating auditory functions.

Interestingly, we did not observe any D1Aa expression in the dopaminergic neurons of the TPp. While a previous study found D1A and D2 mRNA in the TPp of African cichlids (O’Connell et al., 2011), it was not confirmed if D1A was localized to the dopaminergic cells within the TPp. We labeled for the more specific D1Aa subtype in the present study and ultimately the midshipman D1Ab and D1Ba subtype must be tested to confirm the lack of D1 receptors in the dopaminergic cells of the TPp. Lastly, we believe that, by labeling TH-ir TPp neurons coupled with both receptors, we have higher spatial resolution that confirms DAR mRNAs localized to individual cells.

There is currently little to no neuroanatomical evidence for dopamine cells in the TPp receiving dopaminergic input from another brain region. Thus, the robust presence of D2a receptor transcript mRNAs on TPp cells suggests a strong autoregulatory role coupled with presynaptic regulation. In rodents, somatodendritic dopamine release in the ventral tegmental area (VTA) and substantia nigra pars compacta (SNc) has been shown to activate D2 autoreceptors on dopamine neurons, which leads to inhibition via activation of G-protein-coupled inwardly rectifying potassium (GIRK) channels (Aghajanian & Bunney, 1977; Beckstead et al., 2004; Courtney et al., 2012; Lacey et al., 1987; Mercuri et al., 1997). Furthermore, D2 autoreceptors can increase the activity of the dopamine transporter (DAT), thus decreasing dopamine available for postsynaptic signaling (Cass & Gerhardt, 1994; Dickinson et al., 1999; Wu et al., 2002). D2 autoreceptors can also inhibit tyrosine hydroxylase, which is proposed to be a slower, long-lasting mechanism via an inhibition of adenylyl cyclase (AC), which leads to a reduction of cAMP binding to protein kinase A in the regulatory domain of TH (Ford, 2014; Harada et al., 1996; Lindgren et al., 2001; Onali & Olianas, 1989). D2R’s can also serve as pre-synaptic autoreceptors on axon terminals to inhibit the likelihood of vesicular dopamine release (Benoit-Marand et al., 2001; Suaud-Chagny et al., 1991). Altogether, if these functions are largely conserved, the high expression of D2a in these widely-projecting neurons provides support for the regulation of dopaminergic signaling at multiple levels.

An interesting finding from the present study is the variation in relative expression of D2a within individual TH-ir cells in the TPp (see section 3.3, figure 4B). While we did not run a quantitative analysis, it is possible that there is differential D2a expression in various dopaminergic TPp cells that have different downstream targets. For instance, a catecholamine projectome study in larval zebrafish has demonstrated that a certain subset of TPp neurons project to various targets-a small percentage ascend to the ventral telencephalon (e.g., Vv/Vd) while far more project to the hypothalamus and hindbrain/spinal cord (Tay et al., 2011). In midshipman fish, approximately 5-10% of TH-ir TPp neurons project to the inner ear (Forlano et al., 2014). Furthermore, testosterone treatment only affected a subset of TH-ir projections that originate from the TPp, indicating that only some TPp neurons are sensitive to androgens (Perelmuter et al., 2021). While it is unknown if individual dopamine cells in the TPp have extremely widespread projections (e.g. both ascending and descending) in midshipman, the low percentages of TPp neurons that project to specific regions perhaps suggest a more specialized role of each individual TPp neuron. Ultimately, a quantitative tract-tracing study coupled with labeling D2a and TH is needed to determine if TPp cells that project to specific targets differentially express the D2a receptor.

### 4.3 D1Aa and D2a Expression in the Telencephalon

The area ventralis (V) contains nodes that are both auditory recipient as well as part of the vertebrate social behavior network (Goodson, 2005; O’Connell & Hofmann, 2011, 2012). In the present study, we found robust expression of D1Aa and D2a receptors in the Vp and Vv, and only D2a in the Vs. Vp and Vs are proposed to be homologous to the mammalian extended central amygdala/bed nucleus of stria terminalis and the Vv is proposed to be homologous to the septum (Bruce & Braford, 2009; Goodson & Kingsbury, 2013; Maximino et al., 2013; Northcutt, 2008). These regions receive a minority of their input from the TPp and likely undergo complex regulation via dopamine from multiple sources for functions that may not be primarily auditory related (Forlano et al., 2014). Similarly, we observed robust coexpression in both the anterior and posterior parvocellular divisions of the preoptic area (POA), another critical node of the social behavioral network (Goodson, 2005; O’Connell & Hofmann, 2011). Both of these findings support a previous study done in a cichlid fish reporting both D2 and D1a receptor expression in the Vv and POA (O’Connell et al., 2011). Dopaminergic modulation within the POA has been shown to regulate social and reproductive behaviors across vertebrates (Dufour et al., 2005; Hull et al., 1995; Kleitz-Nelson et al., 2010; Nutsch et al., 2016), suggesting that receptor expression in the POA likely reflects a broader integrative role beyond auditory processing. As such, these functions are outside the scope of this paper and will be described elsewhere.

### 4.4 D1Aa and D2a Expression in the Auditory Diencephalon and Midbrain

The TS receives catecholaminergic innervation, likely including both DA (from TPp) and NA (from LC) (Adrio et al., 2002; Hornby & Piekut, 1990), and projects to other higher order auditory processing centers-PGl, AT, and CP (Bass et al., 2000). The TS in midshipman encodes concurrent male vocalizations (Bodnar & Bass, 1997, 1999, 2001) and is thus a critical center for processing social auditory signals. Recent studies have shown excitatory and inhibitory DA modulation of auditory responses in the inferior colliculus (TS homolog) via D2-like receptors in rodents (Gittelman et al., 2013; Hoyt et al., 2019). Furthermore, in line with our findings, D1 and D2 receptor mRNA expression has been localized to the TS in the European eel (Kapsimali et al., 2000; Pasqualini et al., 2009) and the TS, CP, and AT of the African cichlid (O’Connell & Hofmann, 2011). Furthermore, the CP similarly has a diverse response pattern to auditory stimuli, but has broader tuning than neurons in the TS, suggesting that the CP may process more complex auditory stimuli (Lu & Fay, 1995). Altogether, perhaps a robust distribution of both the D1Aa and D2a receptor is required in these auditory regions to efficiently process acoustic stimuli.

The TS also has reciprocal connections to the PAG, which has reciprocal connections with the TPp, CP, and receives input from AT (Kittelberger & Bass, 2013). While the PAG is thought to be a vocal-auditory integration site, its connection to the TPp provides direct anatomical evidence for incoming auditory information modulating the TH-ir cells in the TPp (Forlano et al., 2017; Ghahramani et al., 2018; Petersen et al., 2013; Vetter et al., 2025), which may, in turn, drive downstream changes in dopaminergic modulation of other auditory centers. Interestingly, a study in male midshipman found that dopamine injections in the PAG inhibit vocal production, and this inhibition could only be prevented by combined administration of D1 and D2 antagonists-D1/D2 antagonists administered individually had no effect (Allen et al., 2023). This highlights a unique, biologically necessary functional role for both D1 and D2 receptors within the PAG, and our results colocalizing D1Aa/D2a within individual PAG cells provide an anatomical framework for future mechanistic investigations. The high D1Aa and D2a expression in the PAG may be needed to serve its various roles in vocal-motor and auditory integration. A future comparative study in males and females may elucidate differential receptor patterns and potential functional roles within the various subdivisions of the PAG.

### 4.5 D1Aa and D2a Expression in the Hindbrain

The cholinergic OE is a second nucleus in the brain that projects to the inner ear to modulate auditory sensitivity (Bass et al., 1994; Bleckmann et al., 1991; Tomchik & Lu, 2005). Acetylcholine has an inhibitory effect on inner ear physiology in fish (Furukawa, 1981). Our previous study found a robust summer increase in TH-ir innervation of the OE, and a subsequent tract-tracing study confirmed the source of TH-ir fibers is the dopaminergic cells in the TPp (Forlano, Ghahramani, et al., 2015; Perelmuter & Forlano, 2017). Thus, one would hypothesize that during the summer, reproductive months, higher dopamine innervation of the OE may also be coupled with a high expression of inhibitory dopamine receptors, thus creating further inhibition of the cholinergic OE cells to enhance hearing sensitivity. However, our study reveals that the OE co-expresses both dopamine D1Aa and D2a receptors at roughly the same abundance, although, we have no data on the non-reproductive state. Based on the dense TH-ir innervation of OE neurons, we expected a higher density of D2a and/or D1Aa expression; therefore, it may be that one of the other 5 identified DARs predominates in this nucleus. Notably, the OE in the European eel also expresses both D1 and D2 receptors (Kapsimali et al., 2000; Pasqualini et al., 2009; Roberts et al., 1989). Previously documented seasonal changes, such as increases in BK channels and the density of sensory hair cells of the inner ear (Coffin et al., 2012; Rohmann et al., 2013), likely serve as longer-term structural changes that broadly enhance hearing sensitivity and frequency encoding, while dopaminergic innervation of the OE (and inner ear) allows for a more context-dependent, complex central regulation of hearing at smaller time scales.

The noradrenergic LC receives dopaminergic input from the TPp (Kaslin & Panula, 2001; Ma, 1994), and projects widely to the midbrain and forebrain which likely includes higher auditory areas (Tay et al., 2011). Thus, seasonal changes in CA innervation of these regions may be reflected by a mix of DA and NA. While there are no robust studies investigating NA modulation of central auditory circuits in teleost fishes, studies in rodents and birds have identified NA’s important role in improving the signal to noise ratio of auditory responses and sharpening neuronal tuning in the auditory cortex (Gaucher & Edeline, 2015; Ikeda et al., 2015; Yang et al., 2021). Interestingly, LC cells themselves robustly express both D1Aa and D2a receptors, suggesting that dopamine from the TPp may play a complex role in modulating LC function in addition to its widespread targets. Lastly, the co-expression of D1Aa and D2a in the descending and secondary octaval nuclei similarly suggest a broader pattern in which dopamine from the TPp may provide nuanced, region-specific regulation of auditory processing at multiple levels.

### 4.6 Potential DAR Regulation by Steroids

Previous studies in midshipman have also shown that non-reproductive fish implanted with estrogen or testosterone mimic the peripheral auditory sensitivity of summer, reproductive fish (Sisneros et al., 2004). Furthermore, reproductive females have significantly less D2a receptor expression in the inner ear compared to reproductive females (Perelmuter et al., 2019) and non-reproductive females implanted with testosterone mimic some of the seasonal changes in catecholamine innervation of the auditory hindbrain, as well as the inner ear (Perelmuter et al., 2021). Importantly, androgen and estrogen receptors, along with aromatase (the enzyme that converts testosterone to estrogen) have been localized to the TPp (Forlano et al., 2005, 2010). Thus, steroid regulation of dopaminergic TPp neurons in midshipman is one likely mechanism to modulate auditory sensitivity and processing.

The seasonal surge in gonadal hormones, particularly testosterone, in pre-nesting female midshipman fish (see Sisneros et al., 2004b) may act on androgen receptors on TPp neurons to regulate the expression of D2 autoreceptors, which could contribute to the downstream seasonal reorganization of dopamine innervation of other auditory centers. Testosterone has been shown to influence dopaminergic function and receptor expression in many different organisms. In whiptail lizards, androgen implants in castrated males decrease D1 and D2 receptor expression in the nucleus accumbens (O’Connell et al., 2012). Adolescent syrian hamsters treated with anabolic/androgenic steroids increase dopamine and D2 receptor expression in the anterior hypothalamus (Schwartzer & Melloni, 2010). In adolescent rats, androgens have been shown to increase dopamine transporter (DAT), vesicular monoamine transporter (VMAT), and D2 receptor mRNAs in the substantia nigra as well as decrease D3 mRNAs (Purves-Tyson et al., 2014); furthermore, non-aromatizable dihydrotestosterone mimicked the effect of testosterone, while estradiol did not, suggesting androgen-specific regulatory mechanisms (Purves-Tyson et al., 2014). It is also interesting to note that the effects of androgens on dopamine signaling in rats and hamsters are most robust in adolescence when gonadal hormone levels surge and dopaminergic circuits are still maturing. These findings support the hypothesis that surges in testosterone in pre-nesting female midshipman play a critical role in the robust seasonal reorganization of the dopamine circuit via TPp D2R regulation.

Another proposed mechanism by which steroids regulate dopamine signaling involves estrogen receptor-mediated transcriptional control of DAR expression. Estrogen binds to its receptors, which then dimerize and bind to estrogen response element (ERE), which can serve as transcription factors for specific genes. Previous studies show estrogen treatment in ovariectomized rats increasing D1 and D2 striatal dopamine receptor densities (Chavez et al., 2010; Falardeau & Di Paolo, 1987; Lévesque & Di Paolo, 1991). In non-human primates, D1 and D2 receptor availability fluctuates throughout the menstrual cycle, suggesting that endogenous estrogen levels may dynamically regulate DAR expression (Czoty et al., 2009). Ultimately, while literature supports the hypothesis that estrogen modulates DAR expression, the precise downstream mechanisms remain unclear.

### 4.7 Functional Implications of D1/D2 Coexpression

Robust co-expression of both D1Aa and D2a receptor transcripts suggests the downstream membrane protein receptors are in the same cell. Previous studies have shown that striatal spiny neurons in rodents coexpress D1/D2 (Aizman et al., 2000) and D1A and D2 receptors are coexpressed in Area X neurons in songbirds (Kubikova et al., 2010). Interestingly, dopaminergic neurons that project to mushroom body neurons in fruit flies coexpress the genes that encode D1-like and D2-like receptors and were found at the presynaptic site, suggesting a dual autoreceptor system that precisely modulates dopamine release via D2 inhibition and D1 positive feedback (Hiramatsu et al., 2025). This contrasts with the TPp in midshipman, where we only found D2a; however, the other two D1-like receptors (D1Ab and D1Ba) must be localized to rule out a dual autoregulatory role.

D1 and D2 receptors also heavily differ in dynamics and affinity for dopamine. In vivo, the dopamine affinity of D2R’s has been reported to be ∼100x greater than D1R’s (Beaulieu & Gainetdinov, 2011; Costa & Schoenbaum, 2022; Richfield et al., 1989). Thus, D1R’s require much larger dopamine concentrations for activation compared to D2R’s. Because of the low concentration of dopamine required to activate D2R’s, they are believed to be activated by tonic release of dopamine and only deactivated by transient dips in dopamine whereas D1R’s are believed to be activated by a phasic release of dopamine (Dreyer et al., 2010; Grace, 2000; Missale et al., 1998). Importantly, at a baseline release of dopamine, D2R’s but not D1R’s are proposed to be activated (Costa & Schoenbaum, 2022). We therefore hypothesize that neurons innervated by the TPp (or other DA source) that contain both D1R’s and D2R’s have a baseline level of inhibition through D2R’s and dopaminergic cells in the TPp may either pause firing to release inhibition of the target cell or burst fire to excite the target cell. This multi-layered regulatory mechanism may be necessary for fine tuning neuromodulation and processing complex auditory signals, including discerning reproductive mating calls.

While heavily debated, it has been proposed that, despite D1R’s typically being coupled to Gs proteins while D2R’s are typically coupled to Gi proteins (see section 2), when these receptors are activated together, they can form a hetero-oligomer complex that couples to the Gq protein (Hasbi et al., 2020; Perreault et al., 2011; Rashid et al., 2007). These Gq proteins then activate phospholipase C (PLC), which catalyzes the breakdown of phosphatidylinositol phosphate (PIP) into diacylglycerol (DAG) and inositol trisphosphate (IP3), leading to protein kinase C (PKC) activation and then calcium release (Mizuno & Itoh, 2009). The existence of D1-D2 hetero-oligomers have not come without criticism, however (see Frederick et al., 2015). Considering the robust co-expression of D1Aa and D2a throughout the central auditory system in midshipman, these receptors forming hetero-oligomers are certainly a promising hypothesis that may explain the functional purpose of co-expression. Altogether, these may represent complex mechanisms by which dopamine can act to produce its effects, and further studies are needed to delineate which mechanisms are most relevant for modulating auditory function.

## 5 Conclusions

Here, we provide a comprehensive characterization of dopamine D1Aa and D2a receptor transcripts across the central auditory system of the plainfin midshipman fish, an important model for investigating neural mechanisms of vocal-acoustic communication in vertebrates. The TH-ir neurons of the TPp-the dopaminergic source to various auditory centers that is partly responsible for the seasonal reorganization of catecholamine innervation-exhibits very dense D2a expression while lacking D1Aa, suggesting robust autoregulatory control at both somatodendritic and axon-terminal sites. Nearly all auditory nuclei downstream of the dopaminergic TPp projections contain populations of neurons that either co-express D1Aa and D2a, or express D2a exclusively, suggesting diverse, complex, and region-specific dopaminergic modulation. The previously documented expression of androgen and estrogen receptors within the TPp coupled with prior evidence from teleost and mammalian models demonstrating androgen regulation of dopamine synthesis and receptor expression supports the hypothesis that seasonal fluctuations in gonadal steroids or other seasonally regulated hormones may regulate dopamine receptor expression, particularly within the TPp. However, further studies are needed to confirm the exact mechanism.

Ultimately, these findings reveal a complex dopaminergic cytoarchitecture throughout the midshipman central auditory system. These results lay the neuroanatomical groundwork for future investigations into reproductive state-dependent changes in dopamine receptor expression, functional implications of D1/D2 co-expression, and steroid hormone regulation of dopaminergic signaling pathways. Given the evolutionary conservation of central auditory processing centers across vertebrates that use acoustic communication, these findings provide key comparative insights into dopaminergic modulation of auditory function.

